# Episodic live imaging of cone photoreceptor maturation in GNAT2-EGFP retinal organoids

**DOI:** 10.1101/2023.02.28.530518

**Authors:** Jinlun Bai, David S. Koos, Kayla Stepanian, Zachary Fouladian, Dominic W. H. Shayler, Jennifer G. Aparicio, Scott E. Fraser, Rex A. Moats, David Cobrinik

## Abstract

Fluorescent reporter pluripotent stem cell (PSC) derived retinal organoids are powerful tools to investigate cell type-specific development and disease phenotypes. When combined with live imaging, they enable direct and repeated observation of cell behaviors within a developing retinal tissue. Here, we generated a human cone photoreceptor reporter line by CRISPR/Cas9 genome editing of WTC11-mTagRFPT-LMNB1 human induced pluripotent stem cells (iPSCs) by inserting enhanced green fluorescent protein (EGFP) coding sequences and a 2A self-cleaving peptide at the N-terminus of *Guanine Nucleotide-Binding Protein Subunit Alpha Transducin 2* (*GNAT2*). In retinal organoids generated from these iPSCs, the GNAT2-EGFP allele robustly and exclusively labeled both immature and mature cones starting at culture day 34. Episodic confocal live imaging of hydrogel immobilized retinal organoids allowed tracking of morphological maturation of individual cones for >18 weeks and revealed inner segment accumulation of mitochondria and growth at 12.2 cubic microns per day from day 126 to day 153. Immobilized GNAT2-EGFP cone reporter organoids provide a valuable tool for investigating human cone development and disease.

## INTRODUCTION

Cone photoreceptors are critical for color vision and are impaired in retinal diseases such as cone-rod dystrophy, retinitis pigmentosa, Leber congenital amaurosis, and retinoblastoma (Mustafi et al., 2009; Xu et al., 2014). Modeling of human cone development and disease in animals is challenged by human-specific cone development features (Singh et al., 2018). Human pluripotent stem cell (hPSC) derived retinal organoids (ROs) closely recapitulate human retinal development with similar developmental timeline, cellular composition and laminated structures (Aparicio et al., 2017; Bell et al., 2020). Combined with CRISPR gene editing, ROs provide opportunities to build cone disease models for mechanistic studies and therapeutic screening and are a potential source for retinal cell or tissue transplantation (Gasparini et al., 2019).

hPSC-derived ROs with cell type-specific fluorescent reporters may be used to monitor a cell’s normal and disease-related behaviors. Photoreceptor reporter lines have been generated by introducing a *CRX-*GFP cassette (Kaewkhaw et al., 2015) or a mouse Crx-mCherry cassette (Gagliardi et al., 2018) into the *AAVS1* locus. A rod reporter line was made by replacing the *NRL* coding sequence with EGFP (Phillips et al., 2018); cone reporter lines were produced by inserting a mouse cone-arrestin (mCar)-GFP cassette (Gasparini et al., 2022) or inserting T2A-mCherry at the C terminus of *GNGT2* (Nazlamova et al., 2022); and a retinal ganglion cell (RGC) reporter line was produced by inserting a P2A-tdTomato-P2A-Thy1.2 cassette at the C terminus of *BRN3B* (Sluch et al., 2017). Additional lines reporting multiple cell types include a *SIX6*-GFP / *POU4F2*-tdTomato double reporter line that separately labels all retinal cells and RGCs and a *VSX2*-Cerulean / *BRN3b*-EGFP / *RCVRN*-mCherry triple reporter line that differentially labels retinal progenitor cells, RGCs, and photoreceptors (Lam et al., 2020; Wahlin et al., 2021). These lines have been used to investigate retinal morphogenesis, improve organoid differentiation, and purify specific retinal cell types for transcriptome profiling and cell transplantation.

To build a platform with which to study cone development and diseases, we sought to generate a human cone reporter iPSC line in which EGFP is specifically expressed in both immature and mature cones with minimal disruption of normal cone development. As a candidate, we identified *GNAT2*, which encodes the cone-specific α-subunit of transducin, a G-protein that couples visual pigment opsin to the cone phototransduction cascade (Morris et al., 1997; Morris & Fong, 1993). Prior studies demonstrated that *Gnat2*^*-/-*^ mice exhibit complete loss of cone phototransduction without changes in rod phototransduction or in cone or rod morphology (Ronning et al., 2018). In human retina, *GNAT2* expression is limited to cones, controlled by a cone-specific promoter, and initially induced following the early cone lineage determinants *RXRG* and *THRB* but prior to mature cone markers *ARR3, OPN1SW* or *OPN1LW* (Hoshino et al., 2017; Morris et al., 1997; Welby et al., 2017) (Fig. S1A; data from (Hoshino et al., 2017)). However, unlike *RXRG* or *THRB, GNAT2* does not downregulate in adult cones (Hoshino et al., 2017; Welby et al., 2017). A similar onset order was observed in *CRX-*GFP+ photoreceptors in human ROs, with *GNAT2* first detected at day 37 (Kaewkhaw et al., 2015). These features suggested that the endogenous *GNAT2* promoter linked to a fluorescent protein may serve as an ideal cone-specific reporter.

In principle, cell type-specific fluorescent reporter lines enable continuous or episodic live imaging to characterize developmental and disease processes. Episodic long-term imaging of specific RO regions enables repeated monitoring of individual retinal cells, yet embedding beyond the initial six weeks was shown to impair photoreceptor development (Decembrini et al., 2020; Rashidi et al., 2022). Others have immobilized and imaged mature ROs for up to three weeks (Achberger et al., 2019), yet longer-term imaging has not been demonstrated.

In this study, we produced GNAT2-EGFP cone reporter iPSC lines in which cones are robustly, specifically, and innocuously labeled with EGFP. We also established RO hydrogel immobilization and episodic live imaging methods that enable long-term assessment of individual EGFP+ cone morphological changes, inner segment development, and mitochondria localization. This EGFP-GNAT2 cone reporter line, combined with the immobilization and imaging techniques, provides a useful tool to study cone development and disease.

## RESULTS

### GNAT2-EGFP iPSCs

To assess the suitability of a *GNAT2* cone reporter, we compared *GNAT2* expression to that of other potential cone markers in human fetal, adult, and RO scRNA-seq datasets. In a combined human fetal retina, adult retina, and early-stage RO dataset produced via 3’ end-counting, *GNAT2* was mainly detected in cones from adult retina (Fig. S1B-E; data from Lu et al., 2020). In contrast, our deep, full-length single cell RNA-seq analysis of fetal retinal progenitor cells and photoreceptors showed robust and specific *GNAT2* expression in the majority of fetal cones from post-conception week 13 to 19 (Fig. S1F, G; data from Shayler et al., *in revision*), consistent with the onset timing in bulk RNA-seq (Fig. S1A). This sensitivity and specificity compared favorably with other potential cone markers such as *THRB, RXRG, GNGT2, ARR3, OPN1SW, or OPN1LW*, which were either expressed in both cones and rods (*THRB, RXRG, GNGT2*) or expressed in only a subset of cones (*ARR3, OPN1SW, OPN1LW*) (Fig. S1H-O; data from Shayler et al., *in revision*). These patterns were preserved in human ROs, where *GNAT2* was detected at day 37 in CRX-GFP positive cells (Kaewkhaw et al., 2015) and was specific to cones at an initial day 55 time point (Shayler, et al., *in preparation*).

To generate a *GNAT2* cone reporter line, we used CRISPR/Cas9-mediated homologous recombination to insert an EGFP-P2A cassette at the N-terminus of *GNAT2* in the human WTC11 iPSC derived line WTC11-mTagRFPT-LMNB1 (Allen Institute for Cell Science) (Fig. 1A, B). The N-terminal position of the EGFP-P2A cassette is predicted to enable GNAT2 translation with a single proline residue added to the N-terminus (Fig. 1B). WTC11-mTagRFPT-LMNB1 cells and derivatives express an mTagRFPT-Lamin B1 fusion protein to enable live imaging of nuclei together with other fluorescent protein markers.

**Figure 1.**
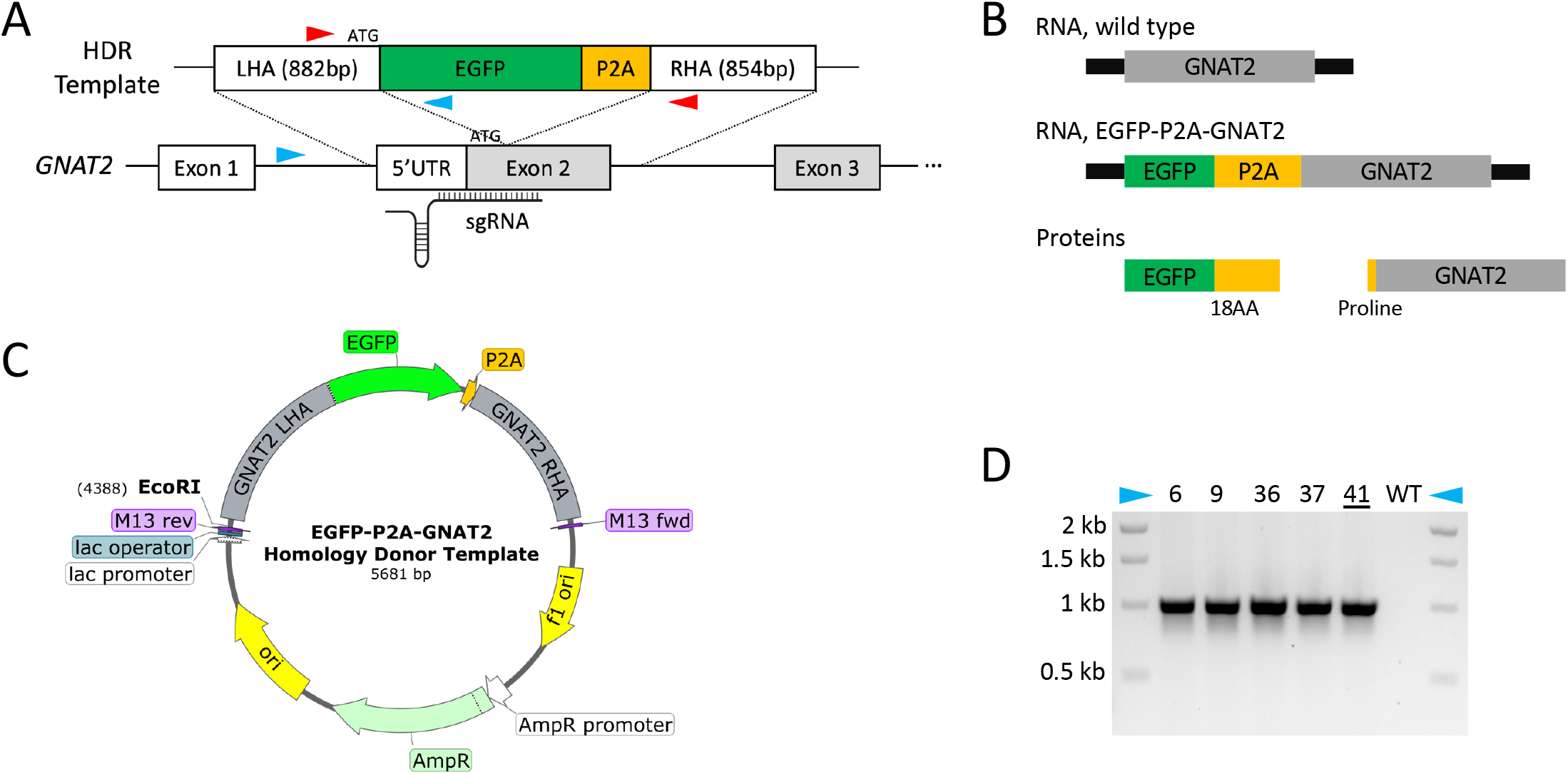
Generation of GNAT2-EGFP iPSCs. (A) Strategy for EGFP-P2A knock-in at the *GNAT2* N-terminus. EGFP-P2A cassette is inserted after the endogenous *GNAT2* ATG start codon. The sgRNA spans knock-in junction. LHA, left homology arm; RHA, right homology arm. Blue arrowheads: location specific genotyping primers. Red arrowheads: insert flanking genotyping primers. (B) Schematic of RNA and protein expressed by wild type and *GNAT2-EGFP* alleles. After translation and P2A cleavage, 18 P2A amino acid residues are added to the C-terminus of EGFP while a proline residue is added to the N-terminus of GNAT2. (C) Homology donor template map with EGFP-P2A cassette flanked by GNAT2 LHA and RHA. (D) Genotyping PCR with insert specific primer pairs. The 1 kb band in C-6, C-9, C-36. C-37, and C-41 indicate knock in of the EGFP-P2A cassette with correct orientation.

Briefly, a homology donor plasmid was constructed by inserting the EGFP-P2A coding sequence between left and right homology arms (LHA and RHA) containing human *GNAT2* genomic sequences 882 bp upstream and 854 bp downstream of the translation start codon (Fig. 1C). The sgRNA spanned the intended insertion site, eliminating the need to introduce a silent mutation on the homology donor plasmid (Fig. 1A). No antibiotic resistance marker was included in the donor vector to enable scarless editing. Following electroporation of the donor plasmid and a plasmid co-expressing *GNAT2* sgRNA and Cas9-T2A-Puro (PX459) (Ran et al., 2013), cells were selected with puromycin, single-cell cloned, and screened by PCR using location-specific and insert-flanking primer pairs (Fig. 1A). PCR with location-specific primers flanking the LHA showed integration of the EGFP-P2A cassette with correct orientation in five of 48 clones tested (Fig. 1D), while insert-flanking primers distinguished two mono-allelic from three bi-allelic knock-in clones (data not shown). The two mono-allelic clones carried mutations on the non-knock-in alleles, whereas the three bi-allelic clones carry no unintended mutations at the knock-in junctions. Partial sequencing of EGFP-P2A-GNAT2 (hereafter referred to as GNAT2-EGFP) clone-41 (C-41) revealed no unintended mutations at any of the top five predicted gRNA off-target sites (Fig. S2A, B; Table S1) and karyotyping revealed no chromosomal abnormalities (Fig. S2C). All subsequent experiments were carried out using this clone.

### GNAT2-EGFP ROs

We next evaluated the GNAT2-EGFP iPSCs’ ability to make ROs with cone-specific EGFP expression. We used a modification of the Kuwahara *et al*. protocol (Kuwahara et al., 2015) to improve RO consistency. Initial culture media was supplemented with small molecule inhibitors of WNT signaling (IWR1) and TGF-β super family signaling (SB431542 and LDN193189) for six days, followed by addition of BMP to induce anterior neural ectoderm and eye field specification (Aparicio et al. in submission). As the parental WTC11-mTagRFPT-LMNB1 had poor early survival, the starting cell number was increased from 12,000 to 48,000, the retinal pigment epithelium induction-reversal was optimally timed from d23 – d28, and long-term maintenance with retinoic acid was begun at d72 (Fig. 2A), which promotes photoreceptor maturation and long-term survival (Kelley et al., 1994; Zhong et al., 2014). During the first 30 days, ROs increased in size and adopted a laminated structure indicative of nascent neural retina. Subsequently, RO growth slowed and ROs shed cells or debris between d80 and d100. A brush border likely representing photoreceptor inner and/or outer segments was evident by ∼ d140 and remained visible until the latest analysis on d245 (Fig. 2B). In mature ROs, EGFP+ cells formed uneven patches occupying the outer-most layer (Fig 2B).

**Figure 2.**
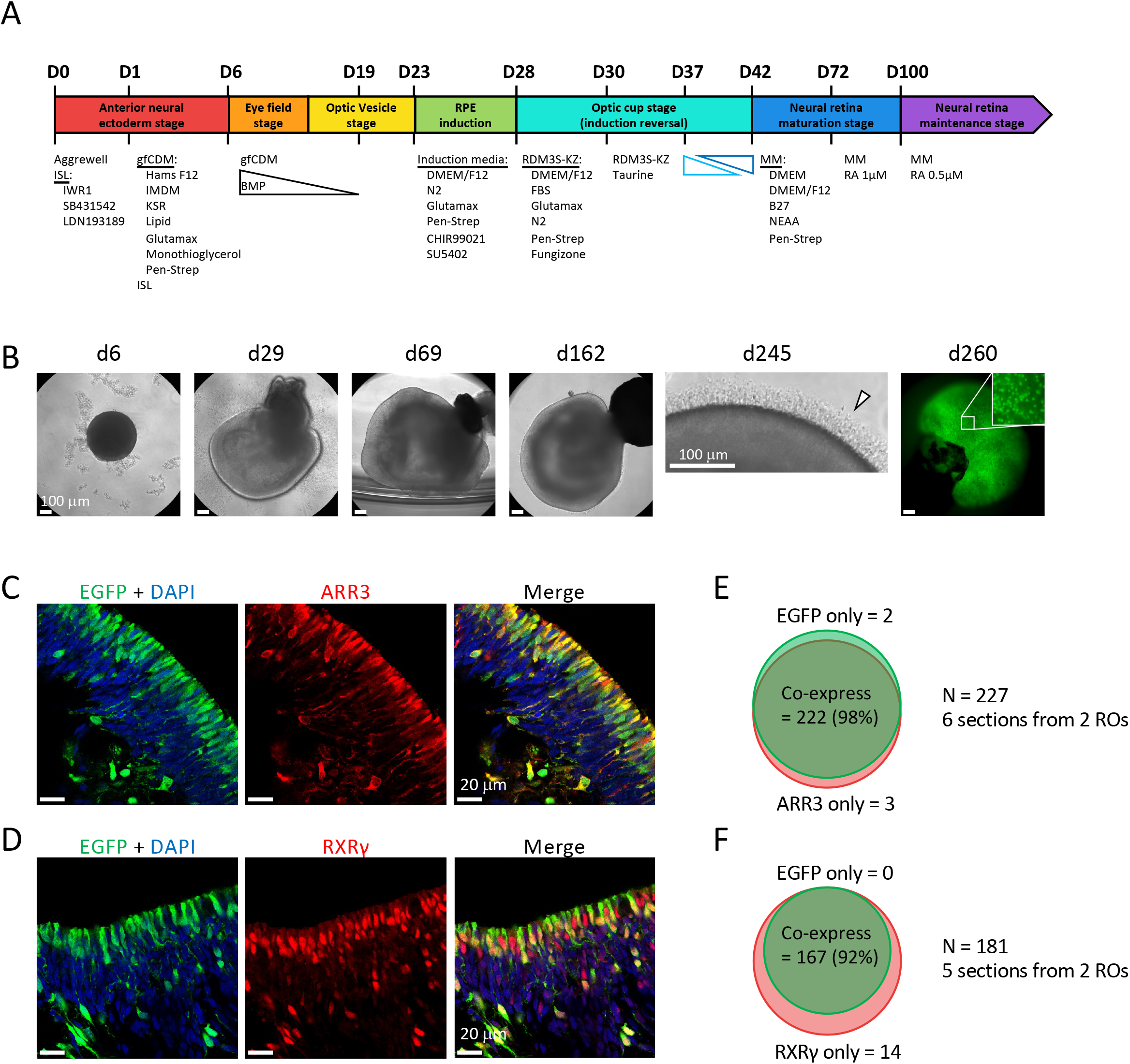
Generation and characterization of GNAT2-EGFP ROs. (A) Overview of RO differentiation protocol. (B) Representative phase contrast images of GNAT2-EGFP ROs at d6, d29, d69, d162, d245 and fluorescent image at d260. White arrowhead indicates visible brush border on a mature RO. Scale bar: 100 µm. (C, D) Immunohistochemistry of d105 GNAT2-EGFP RO indicating co-expression of EGFP and ARR3 (C) or EGFP and RXRγ (D). Scale bar: 20 µm. (E) Quantification of cells expressing EGFP only, ARR3 only or both in six sections from two ROs. (F) Quantification of cells expressing EGFP only, RXRγ only or both in five sections from two ROs.

To evaluate the specificity of GNAT2-EGFP expression, we immunostained d105 RO sections with cone-specific markers ARR3 and RXRγ and assessed their co-localization with EGFP. At this age, most EGFP+ cells had elongated cell bodies occupying the outer-most layer (Fig. 2C, D). Among 227 cells from six sections and two ROs examined for ARR3 and EGFP, 222 co-expressed both, three expressed ARR3 only, and two expressed EGFP only (Fig. 2C, E). Among 181 cells from five sections and two ROs examined for RXRγ and EGFP, 167 co-expressed both, 14 expressed RXRγ only, and none were EGFP+ and RXRγ negative (Fig. 2D, F). These results are consistent with the sequential onset of RXRγ, GNAT2-EGFP, and ARR3 expression during cone maturation (Welby et al., 2017) and confirm the specific and robust labeling of cones in GNAT2-EGFP ROs.

### High-resolution live confocal imaging of cone maturation

We next assessed the feasibility of high-resolution live imaging of GNAT2-EGFP ROs to monitor development of cone cells over time. We performed episodic live confocal imaging on GNAT2-EGFP ROs and captured Z-stack images at different maturation stages, with cones represented by cytoplasmic + nuclear EGFP and nuclear membranes represented by mTagRFPT-Lamin B1 (Fig. 3A, B). From d62 to d111, EGFP+ cell bodies elongated and gradually populated the outer-most organoid layer. By d147 they developed more mature cone morphology, with the appearance of inner segments and pedicles similar to those in older ROs. By d195, cones retained similar morphology and cell bodies were often displaced away from the outermost layer as previously described (Gasparini et al., 2022). Unexpectedly, the mTagRFPT-Lamin B1 nuclear envelope signal was weaker for cones than other retinal cell types, possibly related to the slow turnover of Lamin B1 protein in photoreceptors (Razafsky et al., 2016).

**Figure 3.**
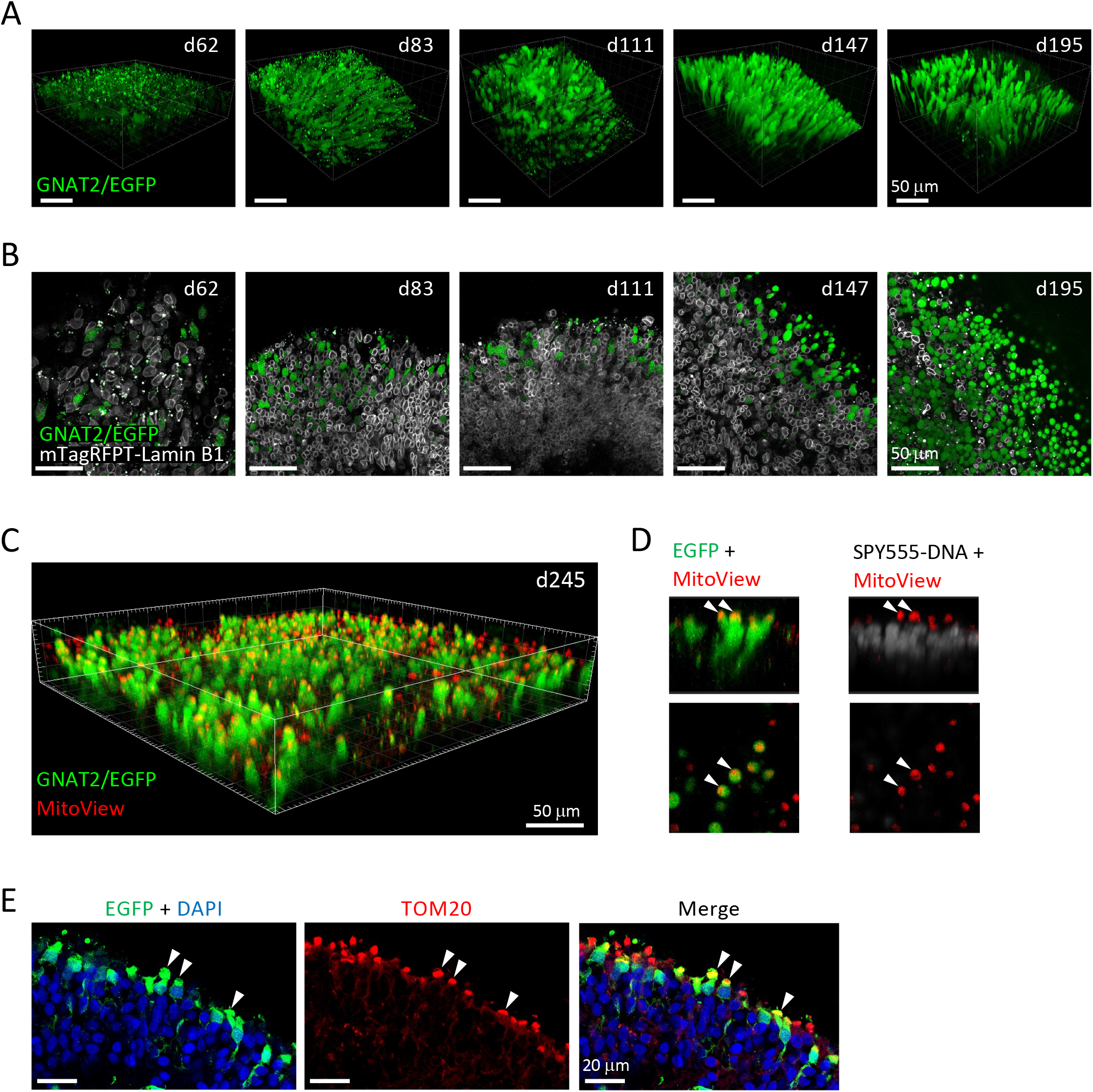
Live confocal imaging of GNAT2-EGFP ROs. (A) Representative 3D renderings of Z-stack live confocal images of GNAT2-EGFP ROs at d62, d83, d111, d147 and d195. EGFP+ cells adopted mature cone morphology in late time points (d147 and d195). Scale bar: 50 µm. (B) Single Z-section from the Z-stack images from (A) in which nuclei are marked by mTagRFPT-Lamin B1 and cones are marked by EGFP. Scale bar: 50 µm. (C) Representative 3D rendering of live confocal Z-stack images of d245 GNAT2-EGFP RO incubated with MitoView. Scale bar: 50 µm. (D) Cross-section images from the Z-stack image in (C) showing coalescence of mitochondria at the EGFP+ cone inner segments above the nuclear layer stained with SPY555 DNA. (E) Immunohistochemistry of d245 GNAT2-EGFP RO confirming coalescence of TOM20+ mitochondria with EGFP+ cone inner segments. Scale bar: 20 µm.

Live confocal imaging also enabled assessment of organelle development. At d245, MitoView staining showed that mitochondria coalesced at the mature cones’ inner segment ellipsoid bodies (Fig. 3C, D), as confirmed by immunostaining with mitochondria marker TOM20 (Fig. 3E).

### Cone maturation in hydrogel immobilized GNAT2-EGFP ROs

To evaluate the maturation of individual cone cells, ROs were embedded in HyStem-C™ hydrogel on Millicell™ cell culture inserts and the same organoid regions repeatedly imaged during long-term culture (Fig. 4A). HyStem-C™ is based on hyaluronic acid and collagen crosslinked polymers and was chosen both because its rigidity can be tuned and because hyaluronic acid and collagen are major components of retinal extra cellular matrix and vitreous humor (Achberger et al., 2019; Hemshekhar et al., 2016; Tram & Swindle-Reilly, 2018) and deemed likely to be biocompatible. Pilot experiments showed 0.25 to 1% HyStem-C™ supported long-term immobilization whereas organoids separated from 2% and 4% hydrogel. ROs embedded in 1% hydrogel at d121 or older could be cultured in this manner for at least 120 more days. The embedded ROs displayed continuous morphological changes with appearance of photoreceptor inner segment protrusions suggesting that embedding does not restrict cone growth and maturation (Fig. 4B, C). Indeed, in organoids embedded on d139, mature cones remained stable and individual cells identifiable until at least d267 (18.3 weeks) (Fig. 4D).

**Figure 4.**
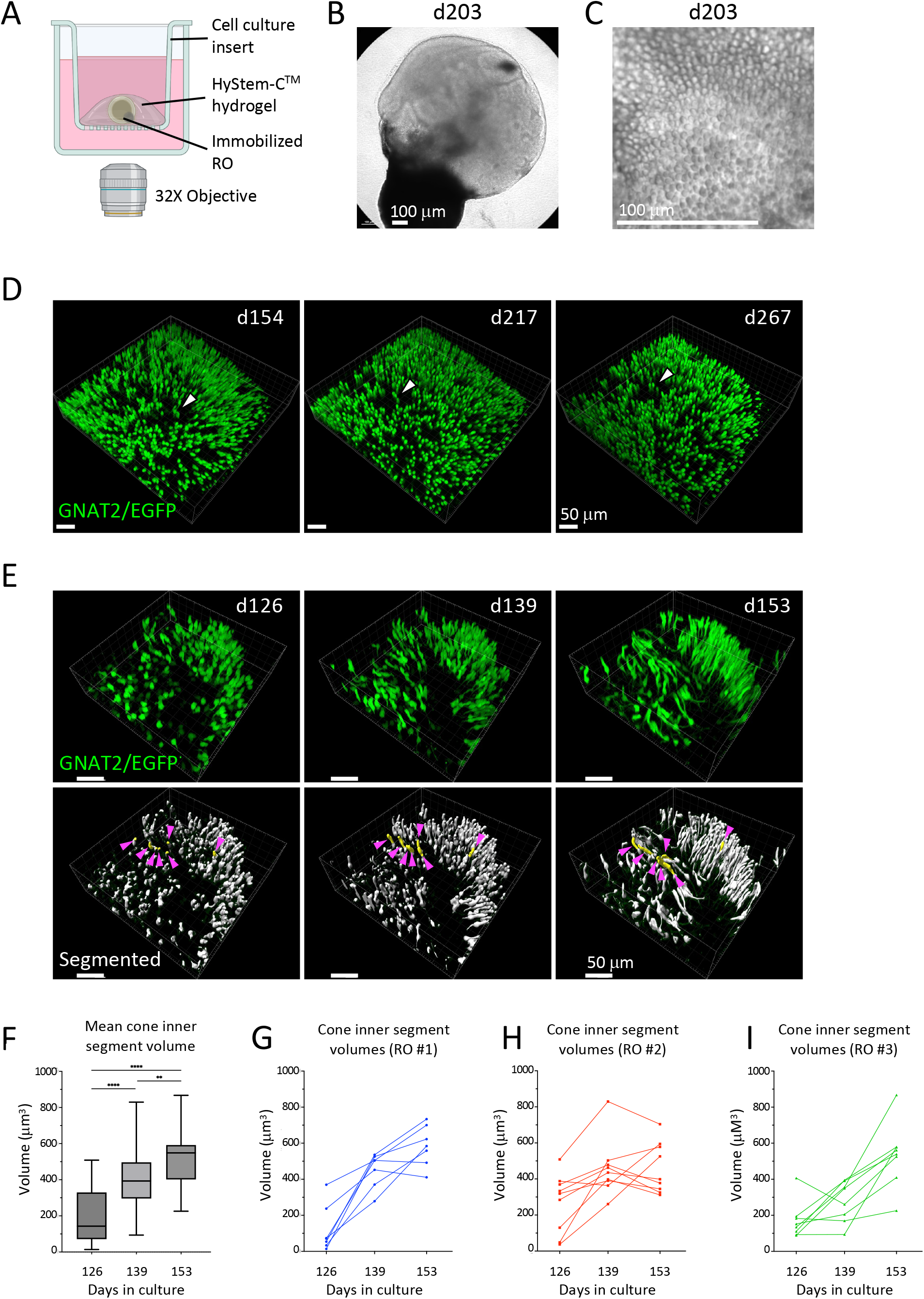
Live confocal imaging of hydrogel embedded GNAT2-EGFP ROs. (A) Schematic of RO embedding and imaging in hydrogel on Millicell™ cell culture insert (created with BioRender.com). ROs are submerged under media and imaged through the bottom of the dish. (B) Phase-contrast image of a d203 RO embedded in hydrogel. Scale bar: 100 µm. (C) Phase-contrast image of a hydrogel embedded d203 RO showing visible photoreceptor inner segment protrusion on the RO surface. Scale bar: 100 µm. (D) Representative 3D renderings of Z-stack live confocal images of hydrogel embedded GNAT2-EGFP ROs at d154 (15 days in hydrogel), d217 (78 days in hydrogel) and d267 (128 days in hydrogel). White arrowhead indicates the same group of three cones at each imaging time point. Scale bar: 50 µm. (E) Representative 3D renderings of Z-stack live confocal images of hydrogel embedded GNAT2-EGFP ROs at d126, d139 and d153 (upper panel) and the EGFP based segmentation result (lower panel). EGFP+ cone inner segment development was tracked. Magenta arrowheads indicate the seven cones tracked and quantified for RO #1 inner segment growth. Scale bar: 50 µm. (F) Pooled quantification of the inner segment volumetric change from 24 cones on three ROs from d126 to d153. (G-I) Inner segment volumetric change of individual cones from d126 to d153.

Episodic live confocal imaging of immobilized ROs allowed us to capture developmental features of the same cells over time. As an example, we used GFP intensity and cell shape-based segmentation on the acquired 3-dimensional Z-stack images to define the volumetric change of 24 cone inner segments in three ROs immobilized at d125 and imaged between d126 and d153 (Fig. 4E). The individual cells were within a defined region where spatial relationships were maintained. We observed the inner segments enlarged from a mean 193 µm^3^ to 523 µm^3^ (p<0.0001) (Fig. 4F), with an average increment of 12.2 µm^3^ per day. However, the initial inner segment size and rate of individual cone inner segment growth varied depending on the organoid and/or organoid region, ranging from a maximum rate of 27.14 µm^3^ per day for a cone from RO #3 to a slight decline of 0.3 µm^3^ per day for one cone from RO #2 (Fig. 4G-I). Taken together, episodic live imaging of hydrogel embedded GNAT2-EGFP ROs enabled long-term evaluation of cone development, such as inner segment morphogenesis and formation of mitochondria-rich ellipsoid bodies, demonstrating the versatility of this reporter system.

## DISCUSSION

In this study, we generated a cone-specific GNAT2-EGFP iPSC reporter line and demonstrated its utility for tracking individual cone development in long-term live-embedded ROs. By tagging *GNAT2* with scarless CRISPR insertion and placing the EGFP-P2A at the N-terminus, we aimed to create a reporter line that faithfully recapitulates cone development with minimal effect on *GNAT2* expression or protein sequence. The GNAT2-EGFP iPSC line robustly labels GNAT2+ cones throughout RO differentiation, with EGFP detected in cone precursor cell bodies as early as d34 and subsequently in maturing cone axon terminals and inner segments. Live imaging revealed that maturing cone precursors develop inner segments and extend pedicles towards the outer plexiform layer between ∼d120 and ∼d150, coinciding with a time of rapid maturation and high glycolytic activity in the photoreceptor layer (Browne et al., 2017).

Recently, two other cone reporter human iPSC lines have been described - one generated by piggyBac mediated insertion of GFP under the control of mouse cone-arrestin (mCar) promoter (Gasparini et al., 2022) and the other generated by inserting a T2A-mCherry cassette into the *GNGT2* locus (Nazlamova et al., 2022). Cone-arrestin is first expressed at a later stage of cone maturation than GNAT2, limiting its ability to label immature cones (Hoshino et al., 2017; Welby et al., 2017) (Fig. S1A). Moreover, only ∼80% of cells labeled with the mCar-GFP reporter were ARR3+ whereas >95% were recoverin+ (Gasparini et al., 2022), potentially reflecting some non-cone expression. Our CRISPR-based editing of *GNAT2* is similar to the Nazlamova et al. editing of *GNGT2*, but used an antibiotic-free scarless knock-in approach so the inserted sequence contains only EGFP-P2A. Additionally, 3’ end counting and our full-length scRNA-seq analyses indicate that *GNAT2* RNA is highly cone-specific, whereas *GNGT2* RNA is expressed in both cones and rods and has no discernable cone-rod specific splicing or exon usage (Fig. S1O) (Lu et al., 2020) (Shayler et al., *in revision*). Nevertheless, GNGT2 protein is reported to be specifically expressed in cones of the developing and adult human retinae (Ong et al., 1997) as is mCherry protein in *GNGT2-T2A-mCherry* ROs (Nazlamova et al., 2022), suggesting that post-transcriptional mechanisms enforce cone-specific translation of GNGT2 and the linked mCherry in *GNGT2-T2A-mCherry* ROs, despite similar cone and rod *GNGT2* RNA levels.

Furthermore, we established a protocol with which to track individual cone development. Live-embedding in HyStem-C™ hydrogel enabled long-term immobilized RO cultures and monitoring of cone development with episodic live confocal imaging. Immobilizing the ROs in thin layers of hydrogel allows nutrient diffusion from all sides and allows RO to grow and mature with minimal restriction (Fig. 4A). Cells were tracked for at least 120 days starting at > d120. The hydrogel maintained structural integrity and transparency during the entire period, allowing repeated imaging of the same RO regions. However, live-embedding might be more disruptive in younger < d80 ROs due to the ongoing rapid organoid growth and morphological changes. The biocompatibility of HyStem-C™ hydrogel was evident from the fairly consistent growth of cone inner segments and may relate to its derivation from hyaluronic acid and collagen, which are major components of vitreous humor and retinal extracellular matrix (Achberger et al., 2019; Hemshekhar et al., 2016; Tram & Swindle-Reilly, 2018). Combining this GNAT2-EGFP cone reporter with further CRISPR editing and live imaging provides a powerful tool to study cone development and disease.

## METHODS

### Human iPSC culture

WTC-mTagRFPT-LMNB1 human iPSCs (Allen Institute for Cell Science) were cultured in feeder-free conditions in mTeSR Plus media (Stem Cell Technologies, #100-0276) on Matrigel (Corning, #354277) coated 35mm dishes. Cells were seeded at 25,000 cells per dish, fed daily, and passaged every 5 days at approximately 70% confluency. When passaged, cells were washed with DPBS (Corning, #21-031-CV) and colonies gently lifted by 2 min incubation in ReLeSR (Stem Cell Technologies, #05872) at 37°C. After neutralization with mTeSR Plus, cells were centrifuged for 3 min at 300 x g and seeded at desired densities.

### Generation of EGFP-P2A-GNAT2 homology donor and sgRNA plasmids

The 882 bp left homology arm (LHA) and 854 bp right homology arm (RHA) were PCR amplified from WTC-mTagRFPT-LMNB1 iPSC genomic DNA with CloneAmp HiFi PCR Premix (Takara #639298). The LHA, EGFP-P2A, and RHA were cloned into pUC118 backbone using In-Fusion Snap Assembly (Takara #638949). The sgRNA targeting the GNAT2 start codon (AAGACGGCAAAT<u>ATG</u>GGAAG) was identified using the online crispr.mit.edu sgRNA designing tool and cloned into the PX459 sgRNA Cas9-T2A-Puro expression plasmid (Addgene #62988) (Ran et al., 2013) according to the accompanying Zhang Laboratory Target Sequence Cloning Protocol. The resulting plasmids were sequenced to confirm correct assembly. A full list of cloning primers can be found in Table S2.

### Generation of GNAT2-EGFP iPSCs

WTC-mTagRFPT-LMNB1 human iPSCs were dissociated into single cells with 5 min incubation in Accutase (Life Technologies, #A1110501) at 37°C. 200,000 cells were electroporated with 500 ng PX459 and 1000 ng of homology donor plasmid in 10 µl electroporation buffer R using the Neon Transfection System (Invitrogen) with one pulse of 1400 V and a pulse width of 20 ms. After electroporation, cells were placed in mTeSR Plus supplemented with CloneR (Stem Cell Technologies, #05888) for 36 hours and then selected in 250 ng/ml puromycin for 36 hours. After a recovery phase of 4 days, colonies were dissociated into single cells with Accutase and seeded into 96-well plates at an average density of 0.5 cells per well to ensure colonies are derived from single cells. Colonies were expanded and genotyped using two primer pairs: an LHA flanking primer pair that produces a 990 bp band from clones with correct integration of the insert, and an insert flanking primer pair that produces a single 1131 bp band from bi-allelic knock-in clones, the 1131 bp band and a 351bp from mono-allelic knock-in clones, or a single band at 351bp from wild type clones. The 1131 bp KI band and 351bp WT allele band were gel purified (Qiagen, #28604) and sequenced to check for potential mutations introduced during editing. To check for potential off-target mutations, we PCR amplified ∼1 kb regions spanning the top five program predicted off-target sites (IDT CRISPR-Cas9 gRNA checker) from the edited clone-41 and the unedited WTC-mTagRFPT-LMNB1 iPSCs and aligned the sequences. All PCR utilized CloneAmp HiFi PCR Premix according to the manufacturer’s instructions. The predicted off target sequences and amplifying primers are provided in Table S1 and S2. The GNAT2-EGFP iPSC line was karyotyped by the Department of Pathology and Laboratory Medicine, Children’s Hospital Los Angeles.

### Retinal organoid differentiation

Retinal organoids were generated following steps modified from a previous protocol (Kuwahara, et al., 2015; Aparicio et al., *submitted*). Briefly, iPSCs were dissociated on d0 with Accutase and resuspended in Aggrewell media (Stem Cell Technologies, #05893) supplemented with 20 µM Y-27632 (Cayman Chemical, #10005583) and “ISL” cocktail, which consists of 3 µM IWR1 (Cayman Chemical, #13659), 10 µM SB431542 (Cayman Chemical, #13031), and 0.1 µM LDN193189 (SIGMA, #SML0559). Cells were plated in round-bottom 96-well plates at 48,000 cells per well in 200 µl media to allow aggregate formation. From d1 to d5, media was changed to gfCDM (45% Hams F12 (Thermo Fisher, #11765047), 45% IMDM (Thermo Fisher, #12440046), 10% KSR (Life Technologies, #10828028), 1X Chemically Defined Lipid (Gibco, #11905-031), 0.5X Glutamax (Life Technologies, #35050061), 450 µM Monothioglycerol (Sigma, #M6145), and 1X Penicillin-Streptomycin (Corning, #30-002-CI)) supplemented with ISL. On d6, gfCDM media was supplemented with 5 nM BMP-4 (R&D Systems, #314-BP-050), followed by 1/2 media change on d9 and d12, 3/4 media change on d15 and full media change on d19 and d21 with fresh gfCDM only without BMP. From d23 to d27, media was changed to RPE induction media that consists of DMEM/F12 (Thermo Fisher, #21331020), 1X N2 supplement (Thermo Fisher, #17502048), 1X Glutamax, 1X Penicillin-Streptomycin, 3 µM CHIR99021 (Cayman Chemical, #13122), and 5 µM SU5402 (Cayman Chemical, #13182). Starting at d28, media was changed to RDM3S-KZ, which consists of DMEM/F12, 10% Fetal Bovine Serum (Omega Scientific, #FB-01), 1X Glutamax, 1X N2 supplement, 1X Penicillin-Streptomycin, and 0.5X Fungizone (Omega Scientific, #FG-70). Taurine (Sigma, #T8691) 0.1 mM was added from d30 onward. On d30, ROs were transferred to 48-well cell culture plate pre-coated with HEMA (Sigma, #P3932-25G). From d37 to d42, media was transitioned to RO maintenance media (MM) adapted from Zhong et al., (2014): 2/3 RDM3S-KZ + 1/3 MM d37, 1/3 RDM3S-KZ + 2/3 MM on d40, and all MM on d42. MM consists of equal volume of DMEM (VWR, #54000-305) and DMEM/F12, supplemented with 1X B27 supplement (Thermo Fisher, #12587010), 1X NEAA, 1X Penicillin-Streptomycin, 1X Fungizone, 10% FBS, 0.1 mM Taurine, and 1X Glutamax. ROs were subsequently cultured in MM with 1 µM retinoic acid (Sigma, #R2625) from d72 to d100 and 0.5 µM retinoic acid from d100 onward. Differences from the Aparicio *et al*. protocol (Aparicio et al, *submitted*) included 1) cultures were initiated with 48,000 cells; 2) only BMP-4 was added on d6, no IWR1; 3) induction reversal was initiated on d23 for five days; and 4) RA was first added on d72.

### Immunohistochemistry and quantification

ROs were fixed in 4% paraformaldehyde for 12 min, washed with DPBS 3 times, incubated in 30% sucrose solution overnight at 4°C, embedded in OCT compound, and cryo-sectioned into 20 µm sections. For immunostaining, slides were washed with TBS, blocked for 1h at room temperature (2.5% horse serum, 2.5% donkey serum, 2.5% human serum, 1% BSA, 0.1% Triton X100, and 0.05% Tween 20 in 1X TBS), incubated in primary antibodies at 4°C overnight, followed by TBS wash, secondary antibody incubation at room temperature for 1h, washing, and mounting with Mowiol with anti-fade. Samples were imaged on LSM710 confocal microscope (Zeiss) and processed using Fiji ImageJ. Primary antibodies are in Table S3.

### RO immobilization, live episodic confocal imaging, and image processing

ROs were subjected to live episodic imaging using a LSM780 NLO confocal microscope (Zeiss), with fitted temperature control and CO_2_ chamber. C-Achroplan 32x/0.85 W Korr M27 lens was used to maximize the imaging distance. Non-immobilized ROs were submerged in RO media and imaged in Lab-Tek 8-well chambered coverglass (Fisher Scientific, #12-565-338). For mitochondria live imaging, ROs were incubated with 1X SPY555 vital DNA dye (Spirochrome, #SC201) over night and MitoView 650 (Biotium, #70075) at 200 nM for 30 min prior to imaging. For RO immobilization, individual ROs were live-embedded in 100 µl 1% HyStem-C™ hydrogel (Advanced Biomatrix, #GS312) on Millicell™ 12 mm cell culture insert with 0.4 µm hydrophilic PTFE membrane (Sigma, #PICM01250). The cell culture insert was then submerged in RO media in a 24-well plate, and media was changed following the same protocol as non-immobilized ROs. For live confocal imaging, the rim of the cell culture insert was marked at the 12 and 3 o’clock positions to indicate orientation, and inserts placed in Cellvis 24-well coverglass bottom plates. Ubiquitous autofluorescent debris on the PTFE membrane was used as points of reference for the regions of interest. Collected 3D Z-stack images were processed using Imaris (Oxford Instruments) and individual cone cells segmented using Imaris Surface function. The inner segment was manually segmented from the cell body at the thinnest connecting point and the volume recorded in Imaris. GraphPad Prism was used for statistical test with repeated measure one way ANOVA and to generate graphs.

## ACKNOWLEDGEMENTS

The authors thank Narine Harutyunyan and Andrew Salas of the Stem Cell Analytic Core Facility and G. Esteban Fernandez of the Cellular Imaging Core Facility of The Saban Research Institute of Children’s Hospital Los Angeles for assistance. Microscopy and image processing were performed at the Transitional Biomedical Imaging Laboratory (TBIL) graciously supported by USC and the CHLA Saban Research Institute.

## AUTHOR CONTRIBUTIONS

Conceptualization: J.B., D.C.; Methodology: J.B., D.S.K., K.S., Z.F., J.G.A., D.C.; Investigation: J.B., D.S.K., K.S., Z.F.; Resources: D.W.H.S., S.E.F., R.A.M.; Writing – original draft: J.B., D.C.; Writing – review & editing: J.B., D.S.K., Z.F., D.W.H.S., J.G.A., S.E.F., D.C.; Supervision: S.E.F., R.A.M., D.C.; Project administration: D.C.; Funding acquisition: J.B., D.C.

---

**Figure S1.**
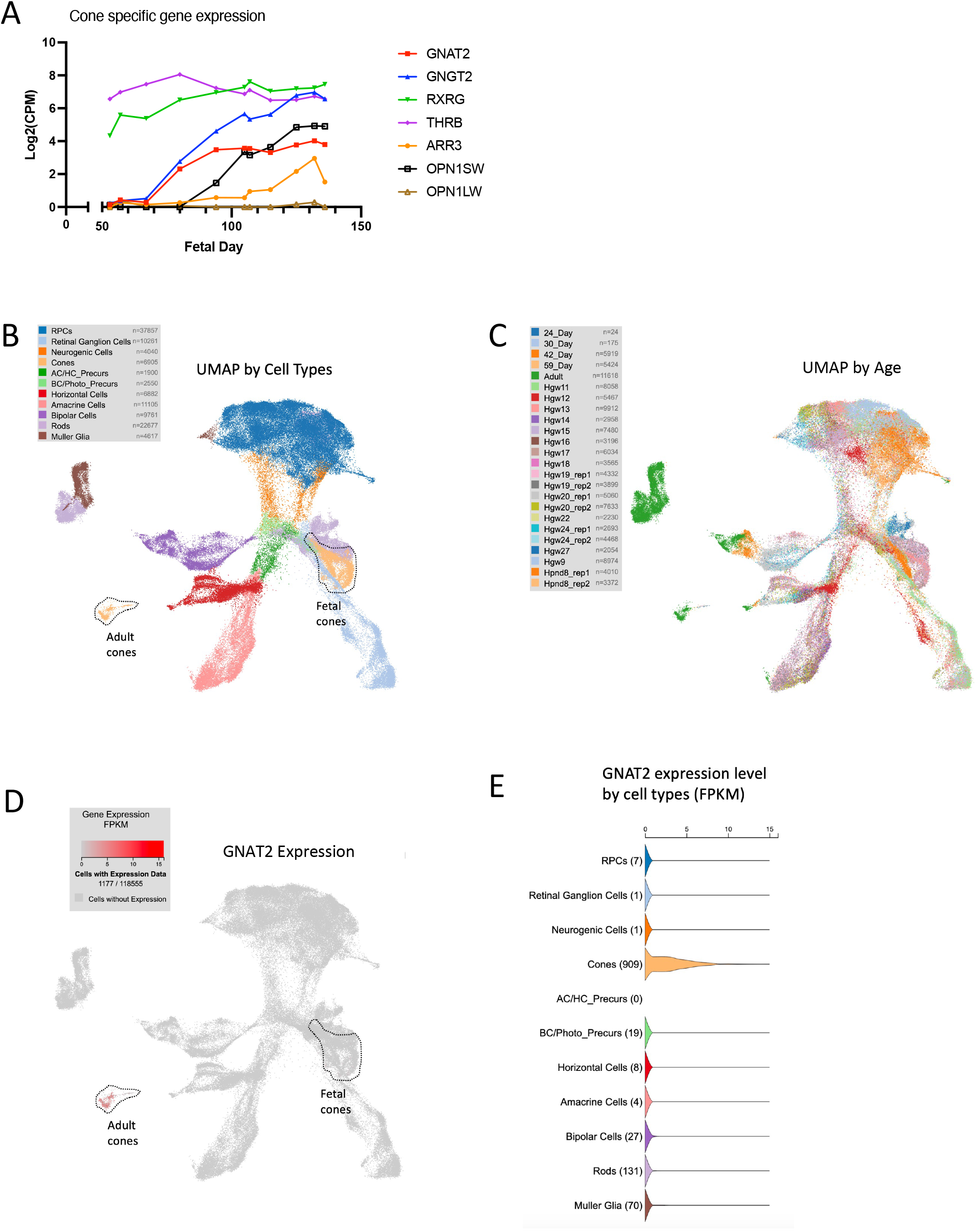
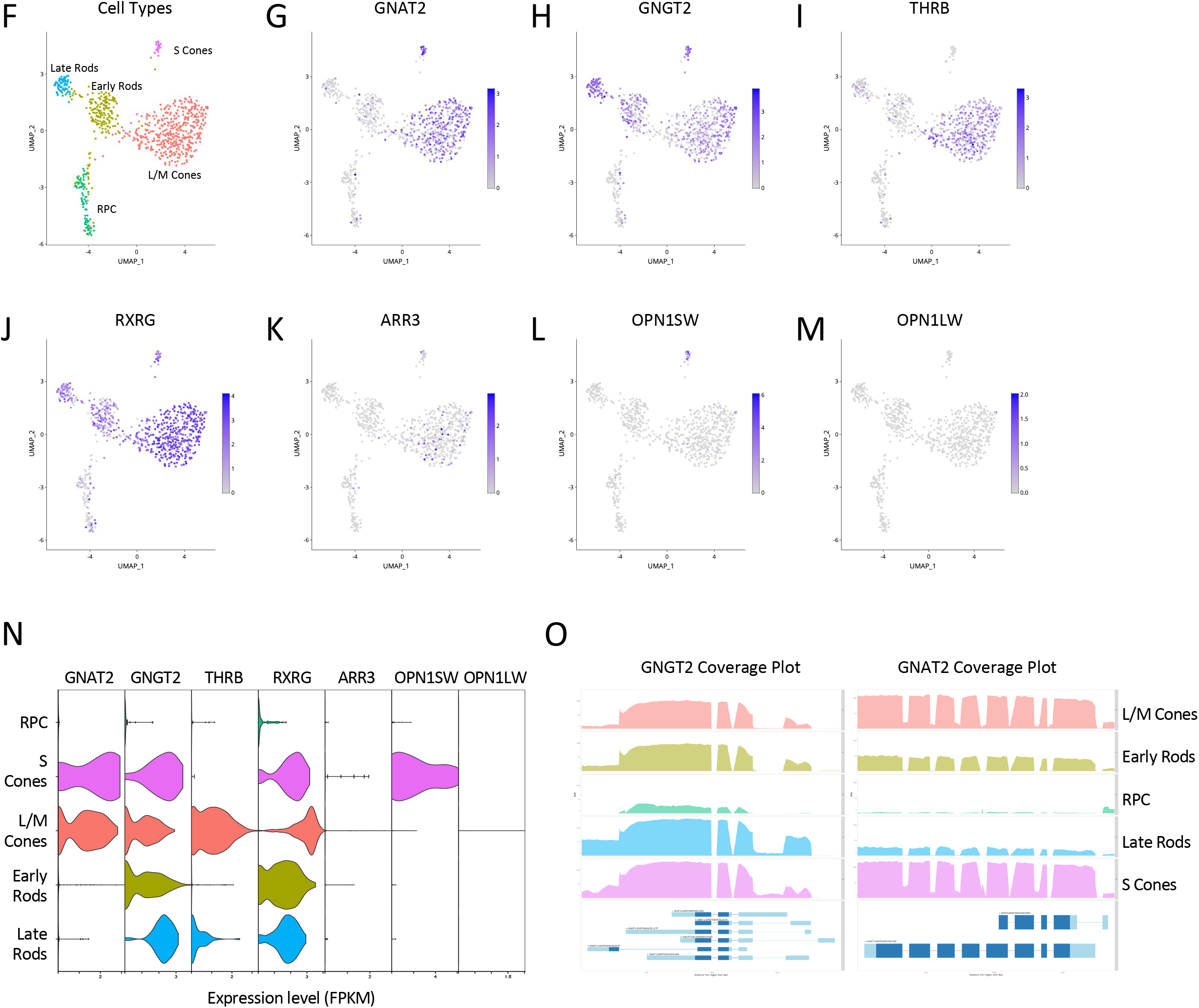
Cone-specific *GNAT2* expression in scRNA-seq analyses. (A) Expression of seven cone markers in developing fetal retina. Bulk RNA-seq data from Hoshino et al., 2017. (B-D) 3D UMAP display of scRNA sequencing of early stage ROs, fetal retinae, and adult retinae from Lu et al., 2020, by cell types (B), tissue age (C) and *GNAT2* expression (D). (E) Violin plot of *GNAT2* expression by cell type from the same scRNA sequencing dataset. (F-L) 2D UMAP display of scRNA sequencing of FACS-enriched fetal retinal progenitor cells (RPCs) and photoreceptors from Shayler et al. (submitted) by cell type (F) and by expression of cone markers *GNAT2* (G), *GNGT2* (H), *THRB* (I), *RXRG* (J), *ARR3* (K), *OPN1SW* (L), and *OPN1LW* (M). (N) Violin plots of cone marker expression by cell type in the same full-length scRNA sequencing dataset. (O) Exon coverage plot of *GNGT2* and *GNAT2* reads by cell type in the same full-length scRNA sequencing. ENSEMBL transcript isoforms are shown below each plot. No cell type specific isoform usage was detected.

**Figure S2.**
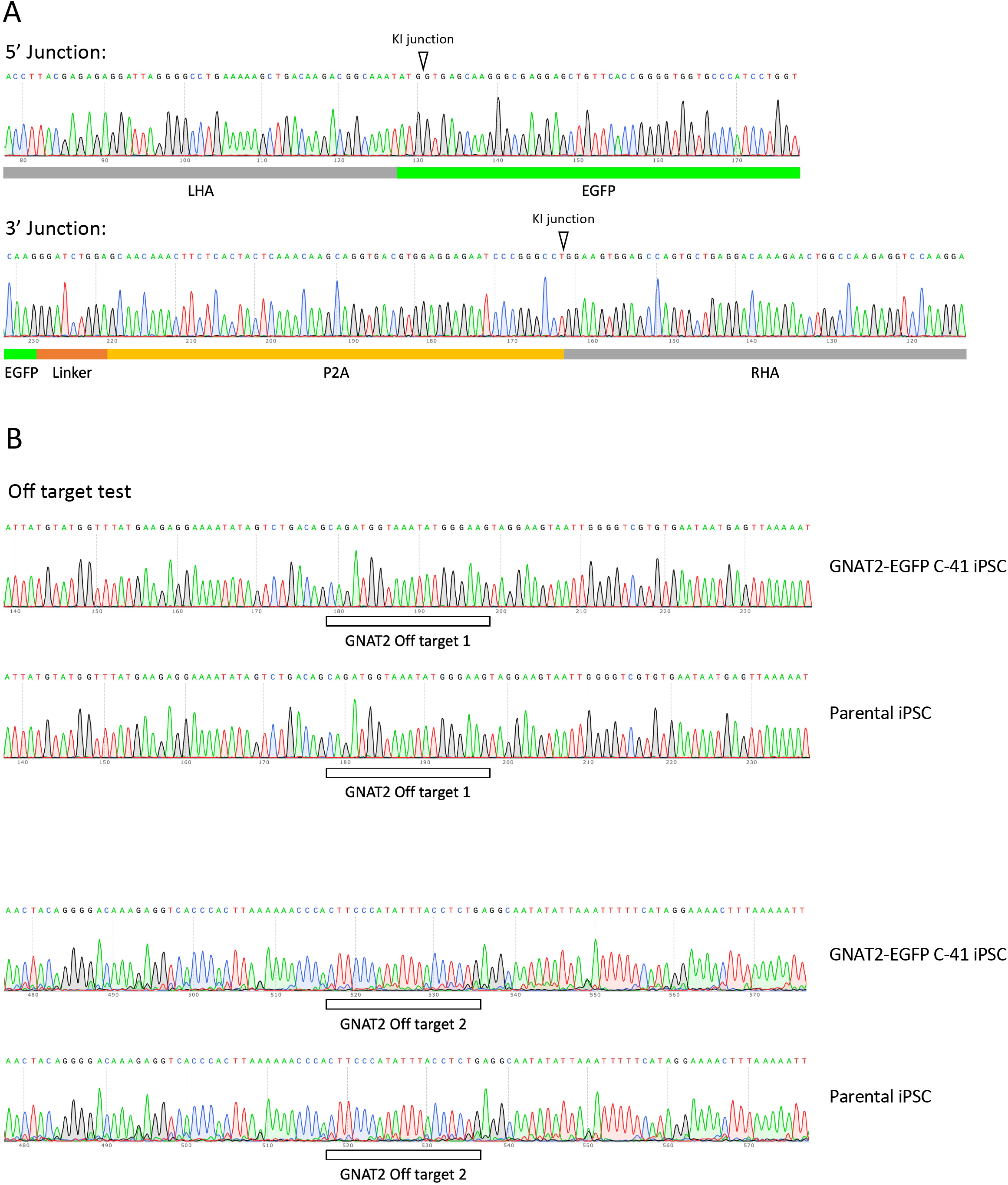
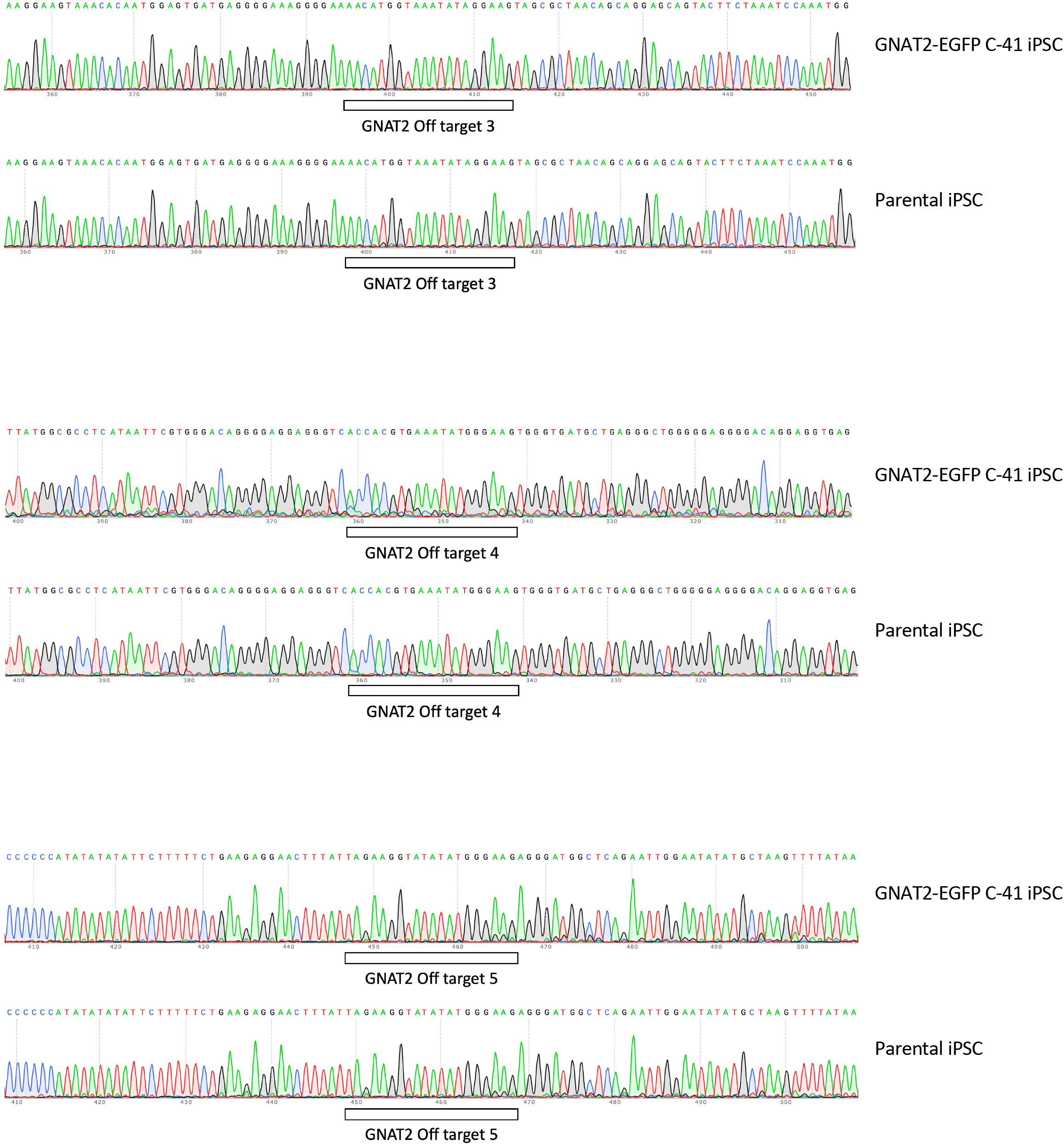
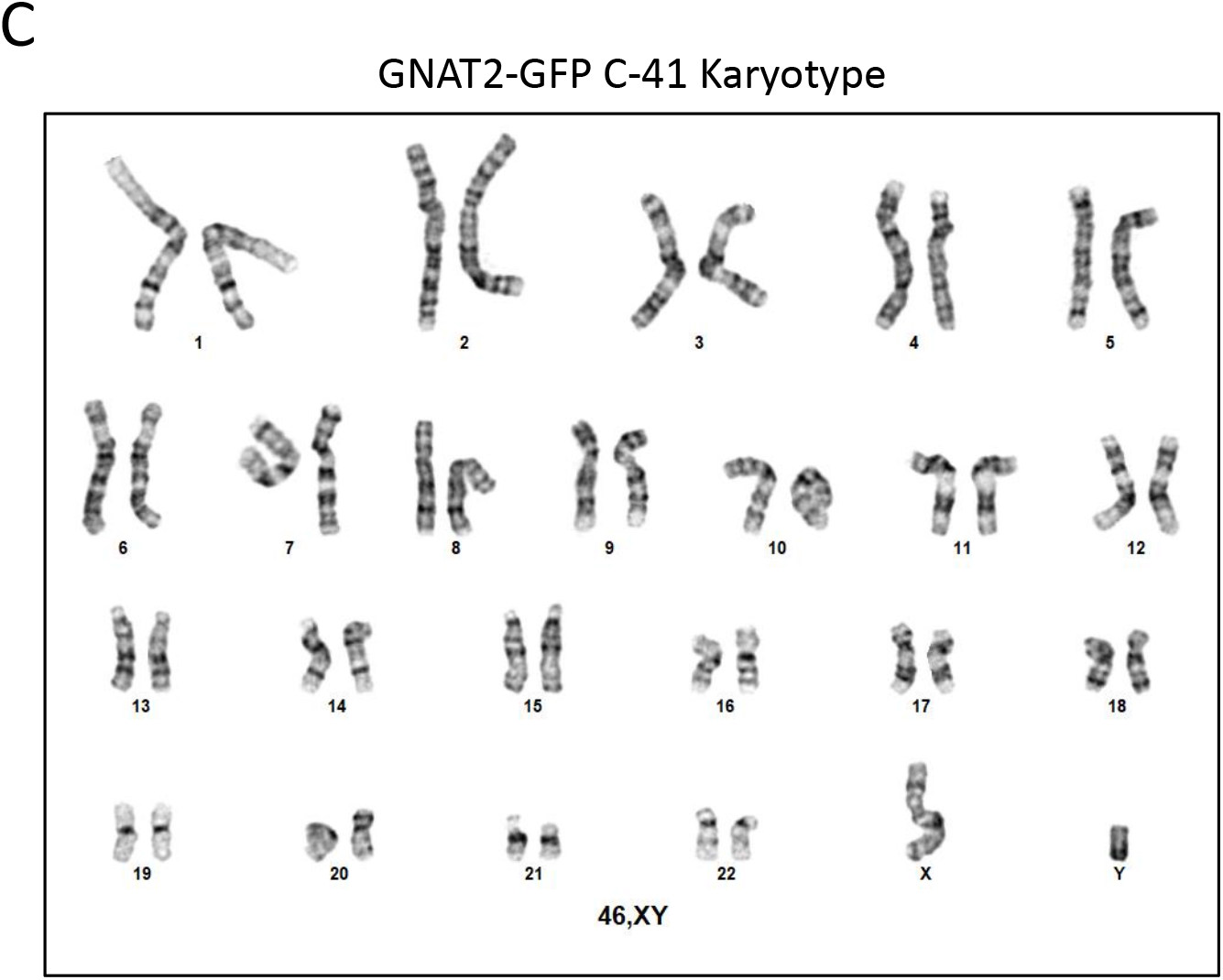
GNAT2-EGFP C-41 genotyping, off target sequencing, and karyotype. (A) Sanger sequencing of the 5’ end and 3’ knock-in junctions showing expected junction sequences. (B) Sanger sequencing showing no mutations detected at the top five predicted off target sites of the gRNA used for CRISPR knock-in. (C) GNAT2-EGFP C-41 shows normal karyotype.

**Table S1.**
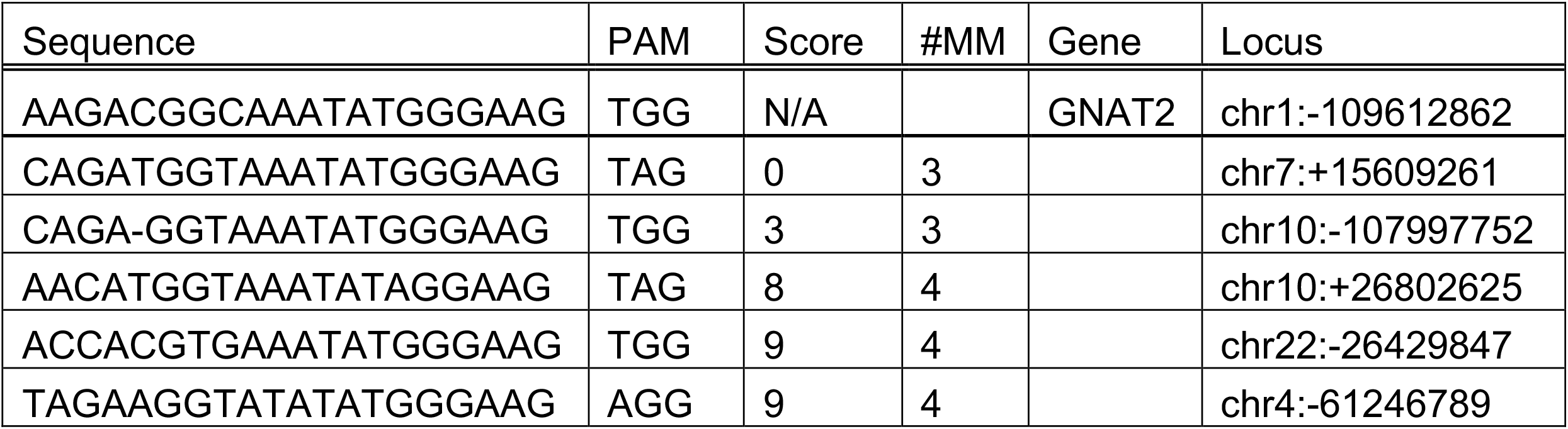
Top five off targets predicted by IDT CRISPR-Cas9 gRNA checker

**Table S2.**
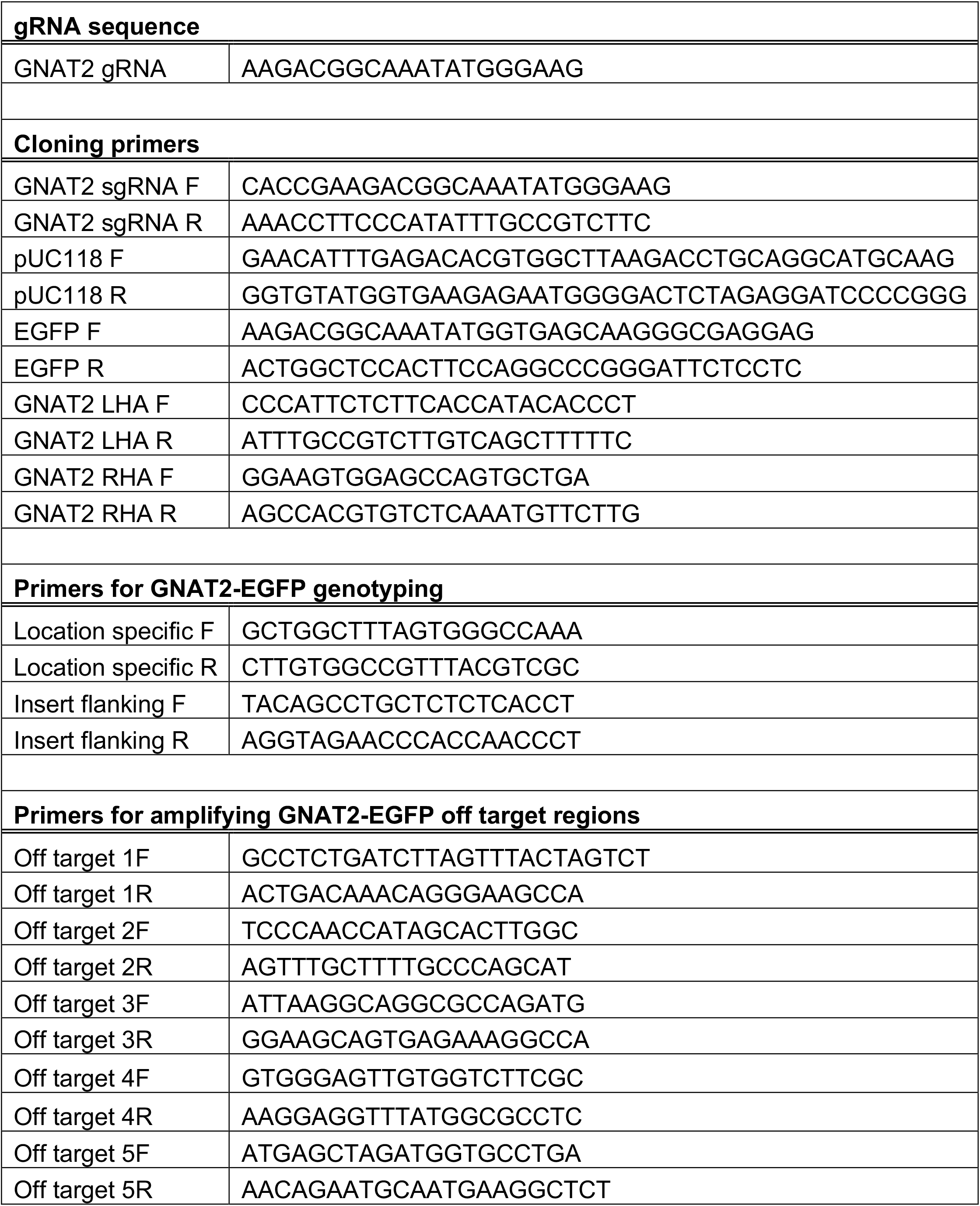
Sequences for primers used in this study

**Table S3.** Summary of primary antibodies used in this study

| **Antibody** | **Supplier company** | **Catalog number** | **Dilution** |
| --- | --- | --- | --- |
| Goat anti GFP | Abcam | ab6673 | 1:400 |
| Mouse anti ARR3 | Millipore Sigma | MABN2636-25UG | 1:500 |
| Rabbit anti RXRγ | Santa Cruz Biotechnology | sc-555 | 1:1000 |
| Rabbit anti TOM20 | Proteintech | 11802-1-AP | 1:200 |

